# Bimodal Coupling Optimization in Biological Rhythms: Balancing Energy Efficiency and Functional Demand

**DOI:** 10.64898/2026.03.25.713094

**Authors:** Jiajun Zhang, Jing Han, Liang-Liang Xie

## Abstract

Biological rhythms are governed by intricate interactions among oscillatory subsystems, yet how they balance functional demands and energy efficiency remains unclear. We present a bimodal coupling optimization strategy where physiological systems dynamically alternate between synchronized (energy-saving) and desynchronized (function-priority) coupling modes. By employing the water-filling principle developed in communications engineering, we prove synchronized heart rate(HR)-respiration oscillations maximize energy efficiency (oxygen uptake per cardiac work). Then, system modeling confirms task/stress-induced oxygen demands enhance oxygen uptake at the cost of desynchronization and reduced efficiency. Experiments reveal a 70.36% decrease in HR-respiration synchronization during arithmetic versus relaxation, enabling 4.43% higher oxygen uptake but with 11.38% lower energy efficiency. This bimodal coupling optimization strategy is also evident in pancreatic islets, with their insulin/glucagon oscillator alternating between in-phase (energy-saving) and anti-phase (rapid glucose reduction) coupling. This framework, integrating engineering and life sciences, reveals a universal regulatory principle for biological oscillatory systems.

## Introduction

Physiological rhythms are pervasive throughout the human body and fundamental to life [1, 2]. Disruptions in rhythmic processes beyond normal ranges or the emergence of abnormal rhythms are associated with diseases [3]. Periodic phenomena are universal, ranging from largescale annual-seasonal cycles to cellular cycles [4, 5]. At the organ level, physiological signals such as heart rate (HR), respiration, blood pressure, blood oxygen saturation, and brain waves exhibit periodic oscillations.

The human body, as a complex system, involves interactions between subsystems, leading to inter-subsystem coupling oscillations (rhythms) across different time and spatial scales: for example, at the organ level, cardiopulmonary interactions coordinate millisecondscale cardiac rhythms with secondscale respiratory cycles [6–8]; at the cell level, circadian clocks, cell cycle progression, and redox oscillations achieve synchronization across hourly timescales through precisely regulated gene expression [9]. This oscillatory coupling is not a static feature but undergoes dynamic reconfiguration with physiological states, such as the restructuring of physiological network topology during sleep cycles [10] and the dynamic adjustment of muscle group coordination patterns under motor tasks [11]. This indicates that the switching of oscillatory coupling patterns embodies system optimization as life wisdom.

However, research on the physiological significance and optimization mechanisms of this dynamic coupled oscillation pattern is limited. Existing theories either focus on single oscillation systems (such as the trade-off between robustness and metabolic efficiency in glycolytic oscillations [12]), with less emphasis on coupling mechanisms; or they address the study of coupling patterns in the context of complex network optimization and regulation, but do not primarily target periodic phenomena [13–15].

We find that the regulatory strategies of biological oscillatory coupling manifest in two key objectives: (1) **maximize energy efficiency** and (2) **optimally meet functional demands**. For example, the synchronization of yeast metabolic gene expression with the respiratory cycle phase optimizes metabolic efficiency [16], and flexible *θ* − *γ* cross-frequency coupling in neural networks can adapt to different tasks [17– 19].

A fundamental question thus arises: does the life system balance energy efficiency and functional demands by strategically switching between two different coupling modes? However, current research on the physiological significance of these coupling modes remains speculative, lacking a rigorous mathematical and mechanistic framework.

In this study, we aim to uncover the optimization principles governing the dynamic regulation of oscillatory coupling in living systems, through investigating the cardiopulmonary coupling system — a critical interaction between two vital subsystems in the human body. The heart is a vital organ that drives blood circulation in the body, sustaining essential life functions [20, 21]. Heartbeat and respiration are two fundamental physiological processes that sustain life. The heartbeat ensures continuous blood circulation, delivering oxygen and nutrients to every cell while removing waste products [22]. Respiration facilitates gas exchange, supplying oxygen to the bloodstream and expelling carbon dioxide [23, 24]. Heart rate and respiration are both periodic oscillations (Fig. 1), and the oscillation of heart rate, specifically observed as the variations in the time intervals between successive heartbeats, is referred to as heart rate variability (HRV). The rhythmic coupling between heartbeat and respiration is vital for maintaining the body’s metabolic balance and energy production. Since S. Hales’ seminal observation of respiration-associated pulse variations in 1733 and C. Ludwig’s groundbreaking kymographic recordings in 1847, cardiopulmonary coupling has remained a classic proposition in systems physiology [25]. It manifests as cardioventilatory coupling, respiratory stroke volume synchronization, and respiratory heart rate variability (RespHRV) [26]. RespHRV[27]—now termed to more accurately reflect its physiological nature than the historical “respiratory sinus arrhythmia” (RSA) [28]—is among the most significant and widely studied of these phenomena. RespHRV is characterized by an increase in heart rate during inspiration and a decrease during expiration (Fig. 1), and as a major contributor to heart rate variability (HRV), it is often quantified via the high-frequency component (HF, 0.15-0.4 Hz)[31], but only when respiration falls within this band.

**Fig. 1:**
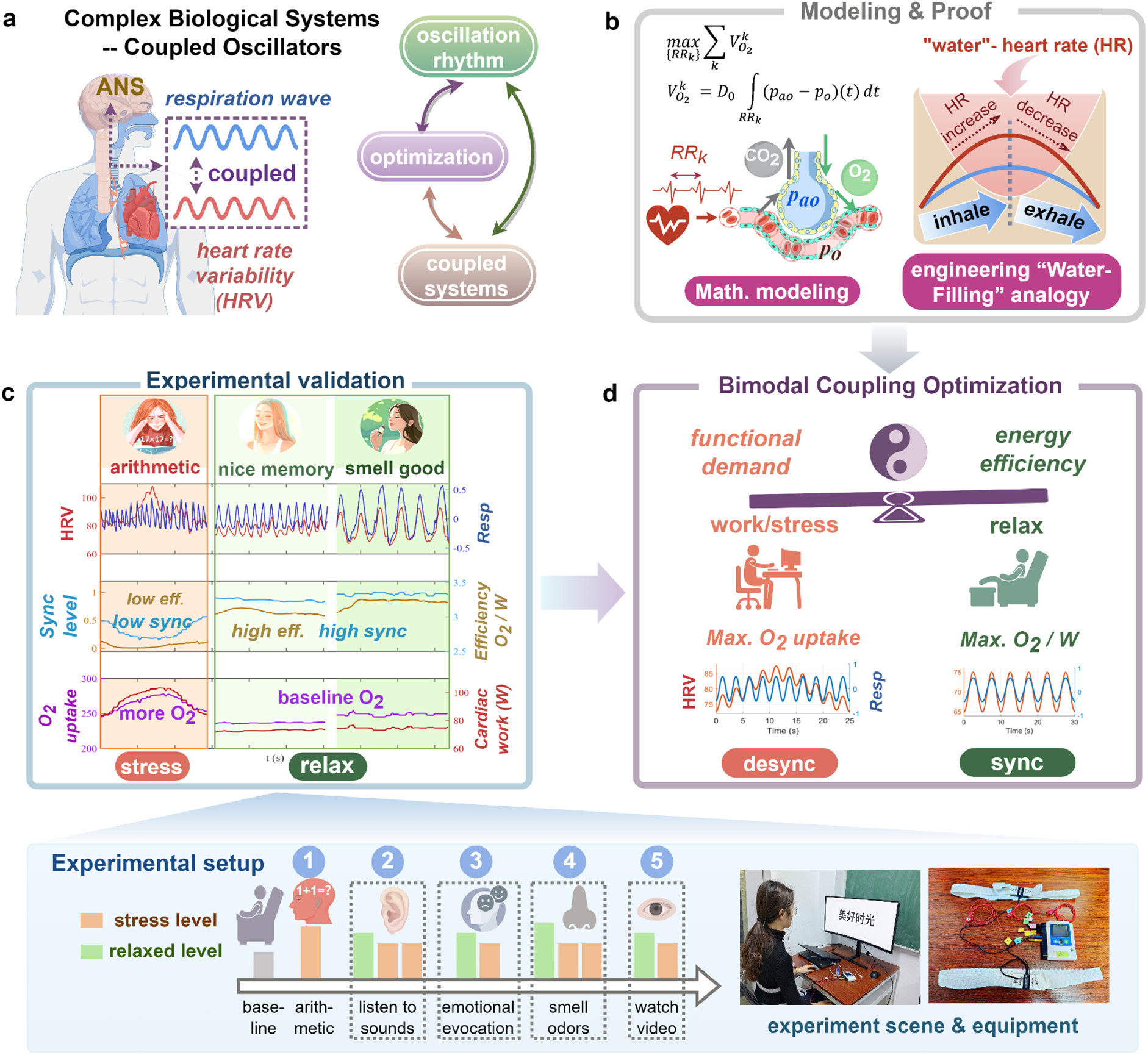
Framework of Bimodal Coupling Optimization in Biological Oscillatory Systems. **(a)** Conceptual overview shows a novel and underexplored intersection of oscillation rhythms, multi-system coupling, and optimization, exemplified by the autonomic nervous system (ANS)-mediated heart-rate variability (HRV) and respiration coupled subsystems. **(b)** Our theoretical approach employs a gas-exchange model together with a “water-filling” analogy (borrowed from communications engineering) to mathematically prove that phase synchronized HR-respiration maximizes energy efficiency (oxygen uptake per unit cardiac work). **(c)** Experimental validation via continuous HR-respiration recordings under three conditions (mental arithmetic, positive memory recall, essential-oil inhalation), with real-time measures of coupling synchrony (*HBC*), oxygen uptake 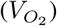, cardiac work (*W* ), and efficiency 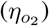. Refer to the bottom image for the complete experimental setup, including the scenario and equipment. The experiment consists of five scenarios, each with different items to induce specific stress/relaxation states. **(d)** summarizes a general bimodal “Yin-Yang” trade-off strategy: desynchronized coupling for “highoutput” (functional demand) and synchronized coupling for “energy-saving” (maximal efficiency), demonstrating nature’s optimal “design logic” and the body’s wisdom. (the figure is drawn by Figdraw, and the photos are of the author.)

Studies have shown that cardiopulmonary coupling enhances the matching of alveolar ventilation and capillary perfusion, thereby improving the efficiency of respiratory gas exchange. Many researches have focused on quantifying the synchronization level of HR-respiration coupling [29–45] to reflect the balance between activities of sympathetic nervous system (SNS) and parasympathetic nervous system (PNS) in the autonomic nervous system (ANS). This is based upon the consensus that when the PNS predominates, RespHRV components is the main contributor to HRV spectrum, whereas SNS activity increases the low-frequency (LF) components of HRV spectrum [31]. Throughout this paper, RespHRV is defined by aligning HRV with the actual respiratory frequency rather than by the fixed HF band. Accordingly, LF denotes the residual, non-respiration-locked component of the low-frequency band, distinct from RespHRV.

Intriguingly, the foundational principle of “heart-breath interdependence” [46] in Daoist cultivation practices also emphasizes the synchronized coupling oscillations. This suggests a profound connection between synchronization, deep relaxation, and energy conservation. Inspired by this, we have developed the Heart-Breath Coherence (*HBC*) metric [29]. *HBC* more accurately quantifies the degree of synchrony between HR and respiration by taking into account dynamic breathing patterns and incorporating the respiration-HR phase difference. This allows for a better assessment of ANS balance. This metric has been demonstrated to significantly enhance performance in assessing the body’s relaxation/stress levels, outperforming the existing 26 HR/respiration-related metrics based on frequency-domain and time-domain analyses [29]. Thus, Daoist wisdom finds resonance in modern science.

The consensus derived from both Eastern wisdom and modern science has raised a critical question: **does HR-respiration synchronization represent an energy-saving mode under the optimization of energy efficiency in the cardiopulmonary system?** Surprisingly, our exploration uncovered profound similarities with the optimal resource allocation strategy based on the “water-filling” principle in communication engineering [47], suggesting the possibility of universal optimization strategies in complex systems.

On the other hand, drawing from the world classic *I Ching* which states that “the interaction of yin and yang constitutes the Dao”, we believe that the work/stress state (corresponding to “yang”) in living organisms also holds physiological significance, just as the relaxed state (corresponding to “yin”) does. Key elements such as HF/LF, PNS/SNS, and relaxation/work can all be viewed as paired “yin” and “yang” aspects. Previous studies have demonstrated that the LF oscillations in HRV spectrum can maintain a robust metabolic level during high-intensity exercise [48]. This prompts us to further explore the physiological significance of the LF components and the physiological optimization logic underlying the dynamic changes in the synchronization of HR-respiratory oscillations.

In this paper, **we propose a theoretical paradigm “*bimodal coupling optimization strategy*” to reveal how the cardiopulmonary system achieves a trade-off between energy efficiency optimization and functional prioritization by switching between high-synchronization and low-synchronization modes**. We quantify the functional demands of the cardiopulmonary system as the volume of oxygen uptake, and its energy efficiency as the oxygen uptake per unit of cardiac work. By integrating a gas exchange dynamic model [49], applying the water-filling optimization theory and numerical optimization approach, we prove that complete synchronization of HR and respiration achieves optimal energy efficiency. Human data from multi-scenario experiments also support this theory: higher HR-respiration synchronization (higher *HBC* values) corresponds to more relaxed state and higher energy efficiency, while maintaining oxygen levels at baseline; under stressful tasks such as mental arithmetic, the demand for oxygen increases, and the rise in heart rate leads to increased cardiac workload. The synchronization between HR and respiration is disrupted (lower *HBC* value), resulting in reduced energy efficiency in the cardiopulmonary system, but oxygen uptake increases. Beyond oxygen uptake, respiration is also vital for the expulsion of carbon dioxide. We also investigate the energy efficiency as characterized by carbon dioxide elimination, and obtained similar results. Fig. 1 vividly illustrates the basic concepts and ideas of this study.

## Results

### HR-respiration synchronization maximizes energy efficiency

We determine the optimal synchronization between heartbeats and respiration to maximize the oxygen uptake per unit of cardiac work. Consider a sequence of heartbeat timings *τ*_0_ < *τ*_1_ < *τ*_2_ < … < *τ*_*n*_. According to the gas exchange model, the total oxygen uptake during the time interval [*τ*_0_, *τ*_*n*_] can be calculated via the following integral (see e.g. [50]):

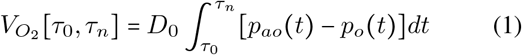

where, *p*_*ao*_(*t*) and *p*_*o*_(*t*) denote the alveolar and blood oxygen pressures respectively, and *D*_0_ = 3.5 × 10^−4^ *L mmHg*^−1^*s*^−1^ denotes the oxygen diffusion constant [51]. Both the variations of *p*_*ao*_ *t* and *p*_*o*_ *t* are not only affected by respiration but also by the heartbeat timings ([49]). Given the time interval [*τ*_0_, *τ*_*n*_], our goal is to determine the optimal timings of the intermediate heartbeats 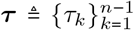 so that the total oxygen uptake (1) is maximized subject to the total cardiac work constraint [50]:

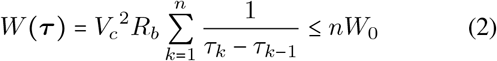

where *W*_0_ denotes the bound on the average cardiac work per heartbeat. We assume that *V*_*c*_ and *R*_*b*_ are constants, following a common modeling choice in the literature [50, 52– 55]. Appendix G discusses cases where *R*_*b*_ and *V*_*c*_ are time-varying.

By solving this optimization problem with the Karush-Kuhn-Tucker (KKT) conditions [56, 57], we have the following result.

#### Theorem 1.

*The optimal heartbeat timings* 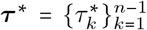 *maximizing* 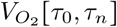 *in (1) subject to the constraint (2) can be determined by solving the following equations:*

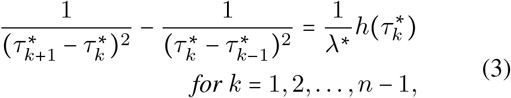

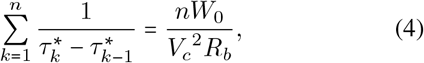

*where the function h(t) depends on the alveolar oxygen pressure p*_*ao*_ (*t*) *and is defined in (16), and the constant λ*^∗^ > 0 *is the Lagrange multiplier associated with the constraint* (2) *and its value can also be determined by solving the equations above*.

Although an explicit analytic solution of (3)-(4) is generally impossible (depending on *p*_*ao*_(*t*), which conversely is affected by ***τ***_∗_), the following property can be easily deduced about the R-R intervals, i.e., the time intervals between consecutive heartbeats.

#### Corollary 1.1

(**Phase dependency of HR on respiration**)

*Consider a single respiratory cycle containing a sequence of heartbeat timings* 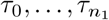, *the optimal R-R intervals* 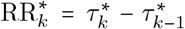 *exhibit respiratory-phase-dependent dynamics:*

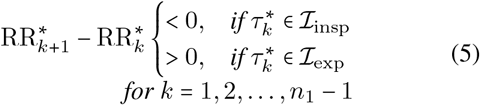

*where* ℐ_insp_ *and* ℐ_exp_ *represent the inspiratory and expiratory phases of the respiratory cycle, respectively. Therefore, as the reciprocal of the R-R intervals, the optimal HR increases during the inspiratory phase and decreases during the expiratory phase*.

Basically, (5) implies that the heart rate speeds up during inspiration and slows down during expiration, which means the phase difference (Δ*θ*) between HR and respiration is 0 ° (as shown in Fig. 2a). Below we elucidate the implications of the theorem and highlight the key ideas underlying its proof. The complete proof is provided in the Methods section.

**Fig. 2:**
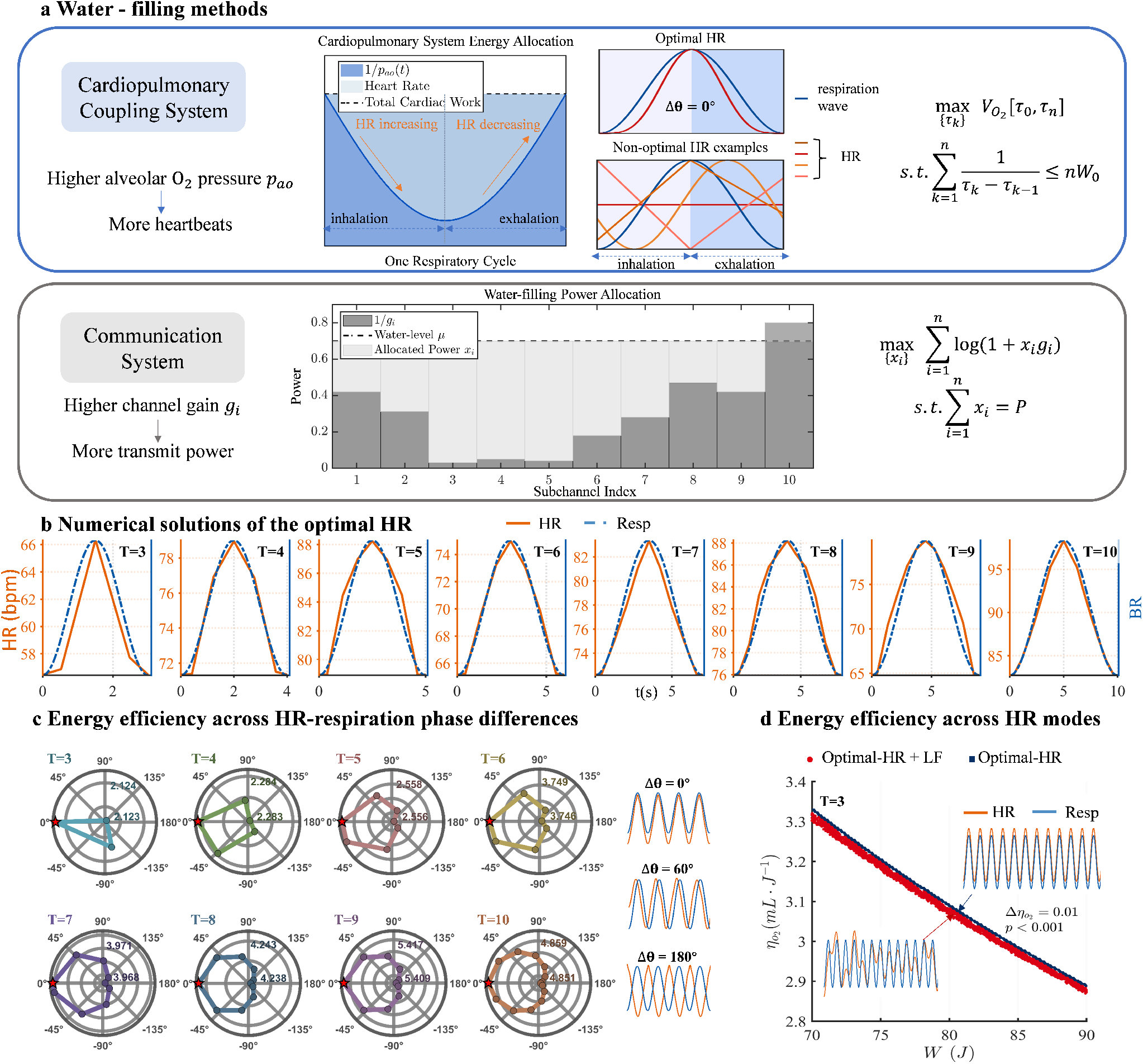
The respiratory-synchronized HR pattern maximizes cardiopulmonary energy efficiency. **(a)** Analogy between the cardiopulmonary optimization problem (blue box) and communication system’s water-filling theory (gray box). The optimal HR pattens show strict phase-dependency with respiration (inspiratory acceleration and expiratory deceleration). The water-filling theory in communication system illustrates channel allocation principles (light gray), and shares similar mathematical foundations between the two optimization problems. **(b)** Numerical solutions of the cardiopulmonary optimization with constrained cardiac work *W*_0_. For given respiratory cycles (T = 3-10 s; blue dotted curves), computed optimal HR (red lines) exhibit near-identical start-to-end values within each respiratory period. It shows that the optimal HR for continuous breathing contains only the respiratory-synchronized component. **(c)** The polar plot visualizes the energy efficiency as a function of HR-respiration phase differences. The distance from the center along each radial spoke represents the magnitude of energy efficiency for that specific phase difference. For respiratory cycles ranging from T = 3-10 s, the chart consistently shows that energy efficiency is maximized when the phase difference is 0 °. **(d)** Energy efficiency comparison between optimal-HR and optimal-HR+LF. For every given cardiac work, respiratory-synchronized HR consistently achieves higher efficiency, further validating the theoretical optimum.

The main idea behind the optimal solution in Theorem 1 is analogous to **the water-filling principle** in communication engineering (see Fig. 2a). Both address constrained resource allocation: the water-filling approach optimizes power allocation across channels to maximize communication capacity, while our cardiopulmonary solution optimizes heartbeat distribution under cardiac work constraint to maximize oxygen uptake. Both adopt a “more for better” strategy—more power for channels with higher gains, and more heartbeats for periods with higher alveolar oxygen pressure.

Within each R-R interval, oxygen uptake mainly depends on the alveolar-capillary partial pressure gradient at the start of the heartbeat. Since blood oxygen partial pressure here equals venous oxygen pressure (≈ 40 mmHg), alveolar oxygen partial pressure primarily governs the oxygen uptake.

This pressure fluctuates with respiration: rising during inspiration, peaking at phase transition, and falling during expiration [58–62]. **Hence, the optimal HR-respiration patterns are characterized by HR increases during inspiration and decreases during expiration, achieving phase synchronization** (see the optimal HR modes in Fig. 2a).

Corollary 1.1 shows that phase-dependent HR—an increase during inspiration followed by a corresponding decrease during expiration—is a necessary condition for maximizing energy efficiency. The optimal heartbeat sequence satisfies the equations (3)-(4) in Theorem 1. Nevertheless, the bidirectional influence between the alveolar oxygen partial pressure and the heart rate renders the analytical solution generally impossible. Instead, we find the numerical solution (details in Methods) for the optimal HR, yielding the results in Fig. 2b.

For the numerical calculation, we choose the cardiac work bound *W*_0_ such that the mean heart rate does not exceed 90 bpm (a physiologically reasonable range supported by experimental data) to facilitate a clearer demonstration of the phase relationship between HR and respiration. These results show that the optimal HR at the beginning of the respiratory cycle is essentially consistent with that at the end. This indicates that the optimal HR pattern for energy efficiency across multiple respiratory cycles should not contain LF components other than the respiratory frequency, meaning that RespHRV accounts for the entirety of HRV spectrum.

To further validate and visualize our findings, we first generate HR-respiration with different phase differences by reordering the optimal R-R interval sequence. As shown in Fig. 2c, optimal energy efficiency is always achieved at phase difference Δ*θ* =0°. Then, we systematically synthesized HR (details in Appendix B) containing varying amplitudes of LF components while maintaining equivalent cardiac work. As demonstrated in Fig. 2c, the energy efficiency 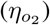 of purely respiratory-synchronized HR (RespHRV-only, blue datapoints) consistently exceed that of LF-containing patterns (red datapoints) across all tested conditions.

Combining the theorem proof and numerical simulations, we demonstrate that HRV with only RespHRV components (i.e., fully respiration-synchronized HR) can achieve maximal oxygen uptake under identical cardiac work constraints.

### Increased oxygen demand requires breaking HR-respiration synchrony

When the human body transitions from a relaxed state to a stressed state, the dominant component in HRV spectrum shifts from RespHRV to LF oscillations [63, 64]. Since HR-respriation synchronization (RespHRV) improves energy efficiency, why does stress disrupt its synchronization? The following analysis shows that the transition from RespHRV to LF as the primary component in HRV spectrum serves to meet functional demands (specifically the increased oxygen uptake requirements of the body).

The analytical approach and logical steps to address this issue are as follows.

1. If a significant increase in oxygen uptake 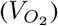 is required, can this be achieved without disrupting the HR-respiration synchronization (i.e., maintaining the same waveform and phase between HR and respiration)? This means that the only variable we can change is the **amplitude of the HR**. Therefore, we first examine whether altering the amplitude of the HR waveform can achieve the goal of significantly increasing 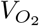 (note that the HR-respiration synchronization and the mean HR must remain unchanged). As shown in Fig. 3, we use sine wave forms to simulate HR and respiratory data under some ideal conditions. Given a respiratory cycle of *T* = 3.7*s* (the average respiratory cycle observed in the mental arithmetic task experiment), the HR patterns are set to be fully synchronized with respiration which can maximize the 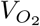. With different mean and amplitudes of HR, the oxygen uptake are calculated by the gas exchange dynamic model [49] (details in Appendix B). Results show that the contribution of only changing the amplitude of the HR to 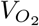 is on the order of 10^−1^ *mL*, which is not sufficient to significantly increase 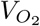 .
2. Since by changing the amplitude of HR does not achieve the goal of significantly increasing 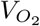, we then attempt to increase the mean HR. Results in Fig. 3 show that increasing the mean HR can significantly increase 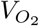. The increase in oxygen uptake is on the order of 10*mL* for every 2 bpm increase in the mean HR, which is significantly higher than the effect of changes in HR amplitude.
3. Since increasing the mean HR is necessary for significantly enhancing 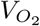, will this inevitably **introduce LF components** in the HRV frequency domain and disrupt the HR-respiration synchronization? Results show that LF components will appear in the HRV spectrum when the mean HR increases or decreases based on a numerical study of a physiological mechanism model of the cardiopulmonary system [65], which is improved and validated by our team [66] (more details in Appendix D). When a person is under stress, the activity of the SNS increases, stimulating an elevation in HR. Therefore, we introduce varying strength of SNS activity, and find that the LF component appear in response to changes in SNS activity (Fig. 4a).

**Fig. 3:**
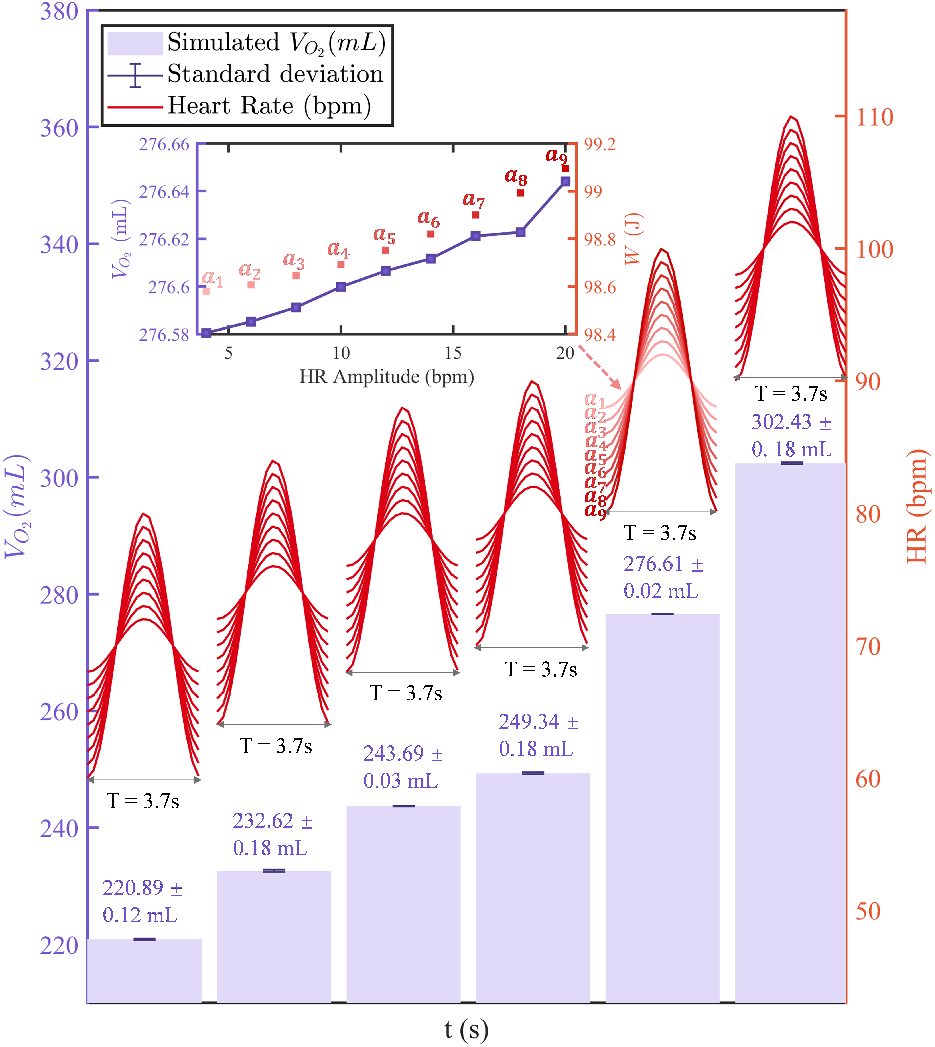
Oxygen uptake 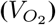 primary depends on mean heart rate (HR) (minimal effect of amplitude modulation). Oxygen uptake 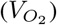 under optimized HR patterns (red curves) across mean HR values (70, 74, 78, 80, 90, 100 bpm) and amplitude ranges (A = [*a*_1_, *a*_2_, …, *a*_9_] = [4,6,…, 20] bpm) at fixed respiratory period (*T* = 3.7*s*). Purple shading and errorbar denote mean values ± standard deviations of 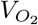 when varying amplitudes at each mean HR. The subfigure shows the changes of oxygen uptake 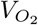 and cardiac work *W* for different HR amplitudes when the mean HR is 90 bpm.

**Fig. 4:**
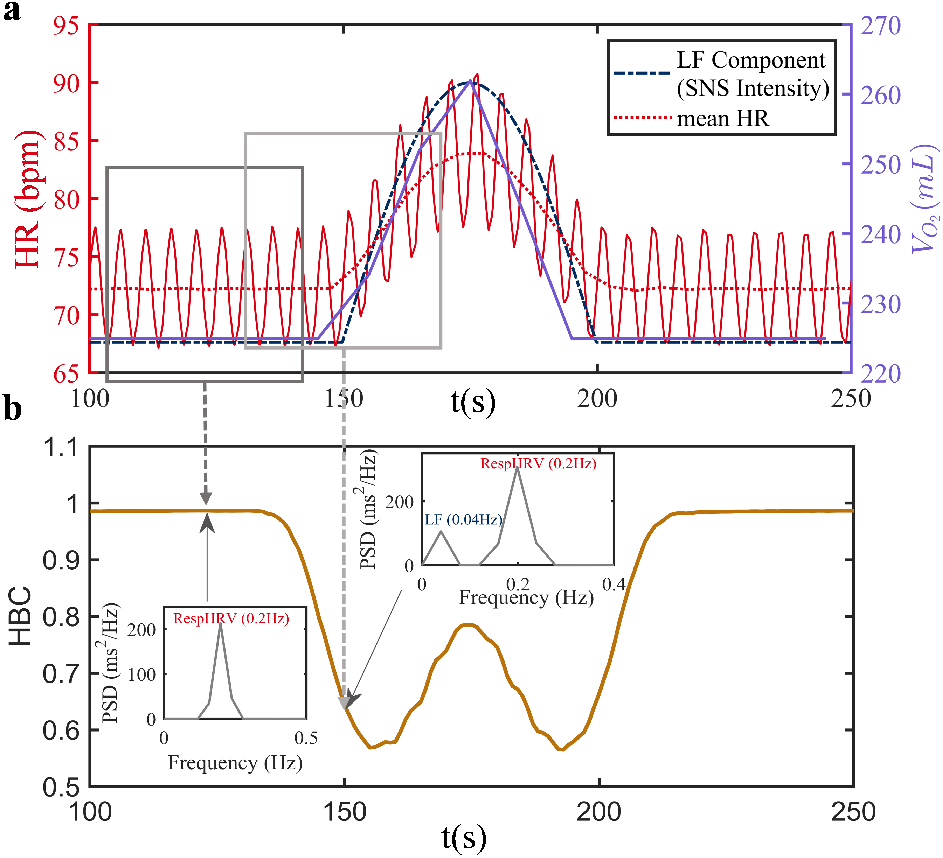
An increase in mean HR introduces LF components and disrupts HR-respiration synchronization, which can be dynamically measured by *HBC*. **(a)** Simulation of HR under sympathetic activation by the cardiopulmonary dynamic model. Blue dashed line, red line, and purple line show sympathetic tone, HR and oxygen uptake respectively. HR increases with enhanced sympathetic input, introducing LF components into HRV spectrum, and oxygen uptake rises accordingly. **(b)** Changes in *HBC* induced by HR in Fig (a). Insets show the power spectral density of HRV. Values of *HBC* decrease/increase as the proportion of LF components in HRV increases/decreases.

These results demonstrate that to increase oxygen uptake, the mean HR must be elevated, which will introduce LF components into the HRV spectrum, disrupt HR-respiration synchronization, and reduce energy efficiency. It means that the LF components in the HRV spectrum achieve functional demands at the cost of cardiopulmonary energy efficiency.

### Bimodal coupling optimization with *HBC*

Now, we have theoretically established the “*bimodal coupling optimization strategy*”: (1) the high HR-respiration synchronization level (RespHRV component) maximizes the energy efficiency; (2) the low HR-respiration synchronization level (LF component) serves to enhance oxygen uptake to meet functional demands.

However, complete synchronization and complete desynchronization modes are rarely observed. Instead, both RespHRV and LF components of HRV spectrum typically coexist simultaneously, with their relative proportions varying.

Therefore, it is necessary to consider a measure of the HR-respiration synchronization level and its relationship with energy efficiency. To address this issue and enable practical application and experimental validation of these theoretical findings with real data, we employ *HBC* to dynamically quantify the ratios of RespHRV and LF components in HRV spectrum (details of calculation of *HBC* are in Appendix C). As shown in the Fig. 4b and Fig. 5, for dynamic HR-respiration curves, *HBC* [29] has been proved that it can well quantify the real-time RespHRV/LF ratio and contain the phase difference between HR and respiration (as the property of optimal HR modes proven in the theorem 1). The HR and respiration here are synthesized in the form of sinusoidal waves (details in Appendix B). When the LF component or the phase difference increase, *HBC* correspondingly decreases. Therefore, we can validate the aforementioned theoretical results by examining the relationship between *HBC* and energy efficiency.

**Fig. 5:**
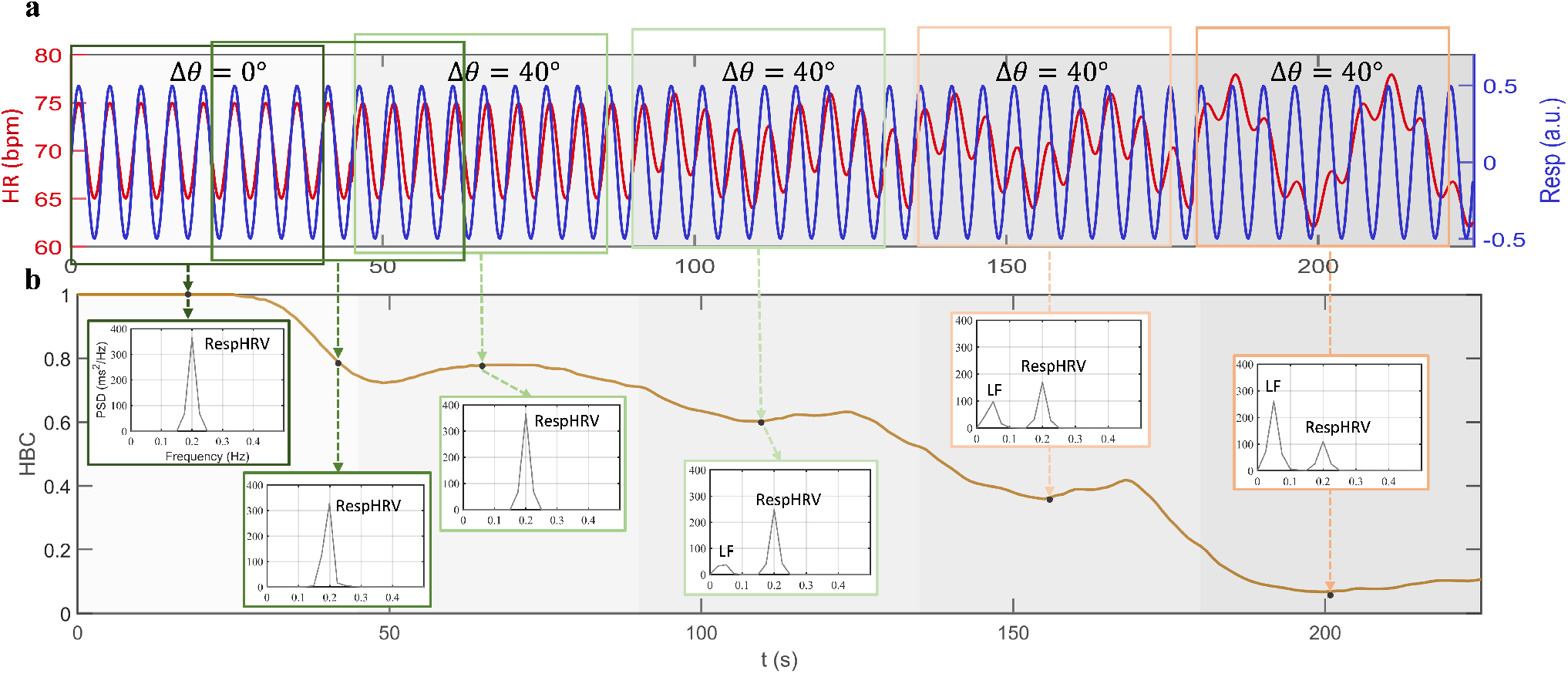
The HR-respiration synchronization level can be dynamically measured by *HBC*. **(a)** Synthetic HR and respiration data with dynamically changing synchronization levels. From left to right, the synchrony of HR-respiration weakens, manifested by increased phase difference (Δ*θ*) and LF components, and a gradual decrease in the proportion of RespHRV components. **(b)** Trends in *HBC* calculated from HR and respiratory data in Fig (a). Insets show the power spectral density of HRV for the corresponding time period. *HBC* values are computed every 1 second, based on a 40-second window of HR and respiratory data centered around the current time point, generating a dynamic curve of HR-respiration synchronization level.

### Experiments show HR-respiratory synchronicity (*HBC*) correlates positively with energy efficiency

Below we validate the “*bimodal coupling optimization strategy*” with experimental data.

We design experiments to induce varying degrees of relaxation and stress in humans, demonstrating different HR-respiration synchronization levels that are effectively quantified by *HBC* in our past research [29]. Specifically, in this paper we have extended the experiments by incorporating a stressful mental arithmetic task and expanding the participant cohort to 65 volunteers. The experimental protocol comprises five scenarios, which contains different items for inducing relaxed or stressed state (Methods for details):

➀ **Arithmetic task** The greatest pressure throughout the entire experiment
➁ **Listening to sounds**, including the blue noise, smooth piano music, and quarrel audio.
➂ **Emotional evocations**, including recalling positive/relaxing moment or annoying moment.
➃ **Smelling odors**, including the essential oil, toilet cleaner spirit, and Chinese herb.
➄ **Watching videos**, including relaxing videos (lovely flower or pet) and a scary spider video.

For each item of the experiment, data of HR, respiration and the fingertip oxygen saturation are measured synchronously, which can be used to derive and calculate the cardiac work (*W*), the volume of oxygen taken up by the blood 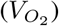, the volume of carbon dioxide output 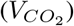, the energy efficiency of cardiopulmonary system (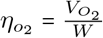 or 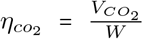) and the HR-respiration synchronization level (*HBC*), and then to verify relationship among them (details in Appendix A). Since the conclusions regarding *CO*_2_ are similar to those for *O*_2_, to maintain focus, subsequent analyses primarily center on oxygen uptake, with certain *CO*_2_-related findings provided in Appendix E.

Since direct measurement of oxygen uptake involves alveolar and blood oxygen partial pressures that are challenging to assess non-invasively and continuously, we employ a gas exchange dynamic model [49] incorporating actual data of HR, respiration for estimation.

The mental arithmetic item represent the most stressful period for volunteers throughout the experiment, characterized by the highest average HR, highest proportion of LF components in HRV spectrum which means SNS is very activie, and the lowest HR-respiration synchronization level (*HBC* value is the lowest). Among the four other scenarios, as reported by volunteers, the most relaxing items are smelling the essential oil, listening to the smooth piano, recalling the positive mement, and watching the relaxed video, during which the RespHRV component is the main component of HRV (higher *HBC* values).

Based on the analysis of the experimental data, here are the findings:

1. Cardiopulmonary energy efficiency (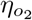 or 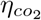 ) is reduced under stress but improves in a relaxed state. The results of 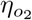 and 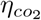 show consistent patterns. Regarding the overall distribution across all volunteers (Fig. 6b), the arithmetic task exhibites significantly lower 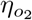 compared to all relaxation-inducing items (Smooth piano *p=* 2.2*e*^−7^, *t* (_47_) = 5.84, confidence interval = [0.17, 0.34], d = 0.74; Positive moment *p* = 6.2*e*^−7^, V = 1039, confidence interval = [0.15, 0.34], d = 0.86; Essential oil *p* 3.4*e*^− 17^, *t* (_61_) = 11.68, confidence interval = [0.47, 0.66], d = 1.48; Relaxed video *p=* 2.1*e*^−8^, *t* (_45_) = 6.78, confidence interval = [0.24, 0.45], d = 1.00). Turning to the analysis of group-level means (Fig. 6e), two key findings emerge: firstly, the arithmetic task yields the lowest mean 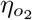 among all experimental items. Secondly, within each of the three other scenarios (emotional evocations, smelling odors, and watching videos), relaxation items consistently demonstrate higher mean 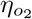 than their stress-inducing items (i.e. for the smelling odors scenario, essencial oil > Chinese herb > toilet). The exception is blue noise in the sound scenario, it may be due to the fact that not all volunteers perceived blue noise as stressful.
2. The energy efficiency (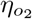 or 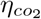 ) demonstrates a statistically significant positive correlation with the HR-respiration synchronization levels (measured by *HBC* values). Firstly, if we examine all items undertaken by each individual, the *HBC* and energy efficiency 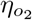 show a positive correlation for 92.2% of the volunteers with mean correction coefficients *r* 0.45 ± 0.16 (Fig. 6d). Secondly, if we examine mean values of *HBC* and 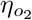 of each item for all volunteers, they show the significant positive correlation, with correlation coefficients of *r =* 0.80 and p=0.0015 (Fig. 6e). Furthermore, values of *HBC* during the arithmetic task for all volunteers are significantly lower than that during other relaxed items which show the same trend as the energy efficiency 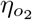 (Fig. 6a and e) (Smooth piano *p =* 0.0011, V = 1442, confidence interval = [0.04, 0.19], d = 0.45; Positive moment *p* = 3.2*e*^−8^, V = 975, confidence interval = [0.08, 0.23], d = 0.50; Essential oil *p= e*^−9^, *t* (_61_) = 9.81, confidence interval = [2.06, 3.12], d = 1.25; Relaxed video *p* = 0.0084, V = 779, confidence interval = [0.03, 0.18], d = 0.51).

**Fig. 6:**
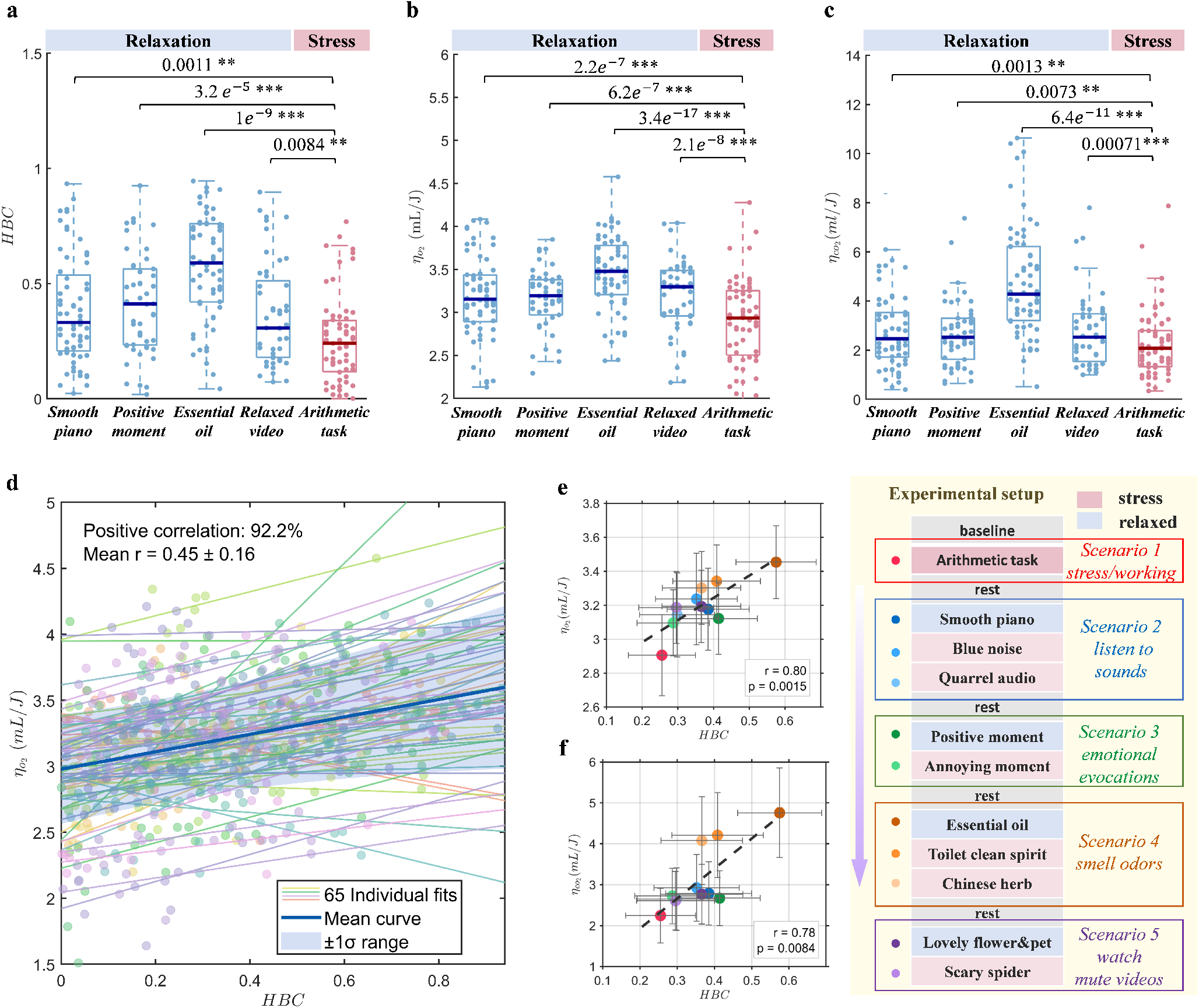
Relaxation enhances the energy efficiency of the cardiopulmonary system (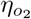 or 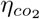 ) which correlates positively with HR-respiration synchronization level assessed by *HBC*. The bottom-right panel shows the experimental setup of five scenarios, each containing items inducing either relaxation or stress. **(a), (b)** and **(c)** respectively show the *HBC*, 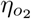, and 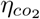 values for 65 volunteers across different experimental items. Numbers above comparison brackets are p-values from pairwise comparisons; see Methods for statistical details. Significance codes: * p < 0.05, ** p < 0.01, *** p < 0.001. During the arithmetic task (a stress state), all three metrics are significantly lower than during other relaxation items. **(d)** Individual and group-averaged 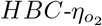 regression lines. Colored points represent data from each volunteer, while lines of the same color show the fitted trend for that individual. The bold dark blue line and shaded area indicate the group mean ± SD. A positive correlation is observed in 92.2% of volunteers (correlation coefficient = 0.45 ± 0.16). **(e), (f)** Mean ± SD of 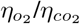 and HBC for each item across all volunteers. The correlation is significant (*p* < 0.01, with *r* = 0.8, *t*_(9)_ = 4.01, confidence interval = [0.48, 1] for 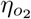 and *r* = 0.78, *t*_(9)_ = 3.80, confidence interval = [0.44, 1] for 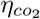).

### Experiments show under stress: cardiac work and *O*_2_ uptake increase

1. Oxygen uptake: during physical or psychological stress, the human body requires more oxygen consumption and uptake. Fig. 7b shows the volume of oxygen taken up during the arithmetic task is significantly more compared to 3 other relaxed items: listening to the smooth piano (*p*<0.001, *t*_(61)_ = -3.84, confidence interval = [-20.42, -6.43], d = -0.49), recalling to the positive moment (*p* < 0.001, *t*_(61)_ = -1.08, confidence interval = [-10.55, 3.17], d = -0.14) and watching the relaxed video (*p* < 0.001, *t* (_45_) = -4.01, confidence interval = [-25.45, -8.43], d = -0.59). The p-value of oxygen uptake between the arithmetic task and smelling to the essential oil scenario is more than 0.05 (V = 854, confidence interval = [-8.95, 3.56], d = -0.14),
2. Cardiac work: obviously, the heart works harder during the working/stress state (Fig. 7a). The overall distribution of cardiac work during the arithmetic task is significantly more than that during relaxed items in each experimental scenario: listening to the smooth piano (*p* < 0.001, V = 294, confidence interval = [-15.08, -6.05], d = -0.71), recalling to the positive moment (*p* < 0.001, V = -78, confidence interval = [-16.46, -6.76], d = -0.88), smelling to the essential oil (*p* < 0.001, V = 74, confidence interval = [- 18.82, -10.41], d = -0.95), and watching the relaxed video (*p* < 0.001, V = 58, confidence interval = [-19.43, -9.83], d = -0.92).

**Fig. 7:**
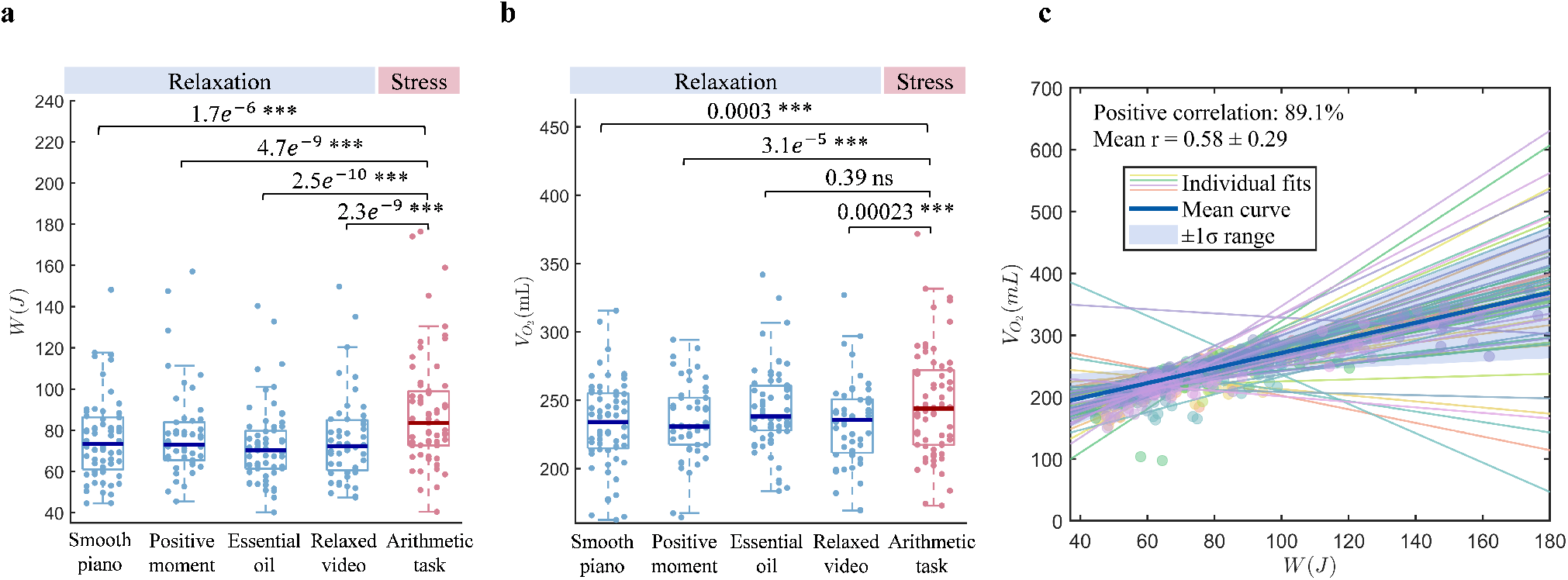
During stress, the cardiac work (*W*) and volume of oxygen uptake 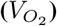 are increasing. **(a)** The cardiac work (*W*) for different experimental items. The *W* during the arithmetic task is significantly more than that during other relaxed items in each scenario. Numbers above comparison brackets are p-values from pairwise comparisons; see Methods for statistical details. Significance codes: * p < 0.05, ** < p 0.01, *** p < 0.001. **(b)** The oxygen uptake 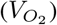 for different experimental items. The 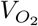 during the arithmetic task is more than that during other relaxed items except the item of essential oil. **(c)** The individual *W* -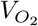 regression line for each volunteer, along with the group-averaged regression line. *W* -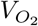 showes a positive correlation in 89.1% of volunteers, with correlation coefficients of 0.58 ± 0.29.

To cope with changes of physical and mental states, the cardiac work and oxygen uptake are changing. Meanwhile, the increase in work done by the heart is often accompanied by an increase in oxygen uptake (Fig. 7c).

Overall, as shown in Table 1, these results confirm that during tasks, the HR-respiration synchronization level and energy efficiency decrease, while the cardiac work and oxygen uptake increase. Specifically, <u>under stress (mental arithmetic task) compared to relaxation, the mean HR-respiration synchronization level decreases by 70.36%, mean energy efficiency decreases by 11.38%, the mean oxygen uptake increases by 4.43%, and the mean cardiac work increases by 15.00%.</u>

**Table 1:**
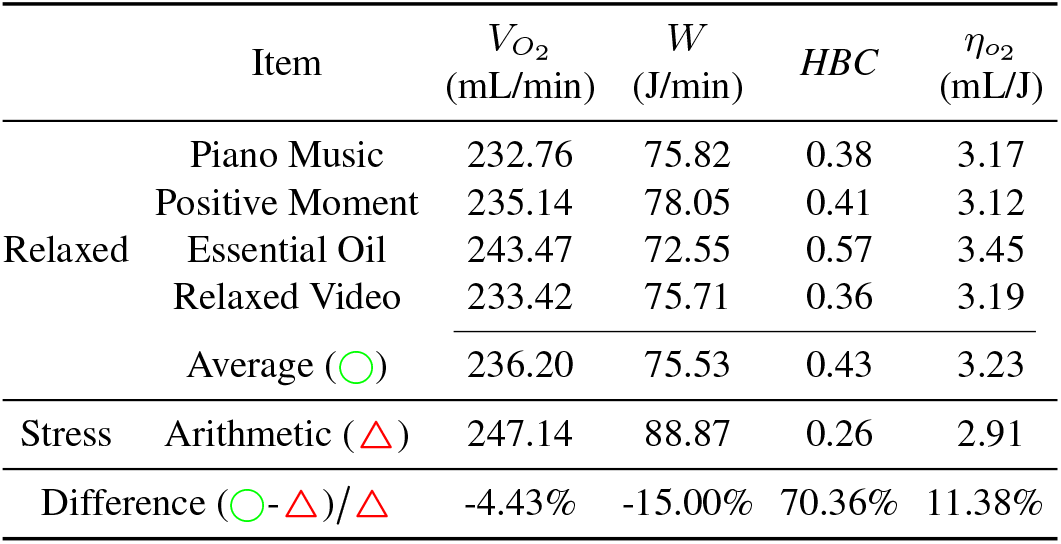
Physiological responses during different tasks.

## Discussion

Through theoretical mathematical proof and simulations based on cardiopulmonary coupling mode and the gas exchange dynamic model, and empirical validation using experimental data, we demonstrate: (1) using water-filling theory, we demonstrate that complete in-phase oscillation between HR and respiration maximizes oxygen uptake while limiting the maximum cardiac work. Through a numerical optimization approach based on simulated annealing, we further confirm that full synchronization between HR and respiration represents the most energy-efficient mode. Experimental data also validate that in more relaxed states, the *HBC*—a measure of HR-respiration synchronization level—increases, leading to enhanced energy efficiency in the cardiopulmonary system; (2) model-based numerical simulations reveal that under high oxygen demand (e.g., during stressful tasks), an increase in mean heart rate (and thus cardiac work) is required to meet demands for oxygen uptake. This introduces LF components into HRV, disrupting HR-respiration synchrony and reducing energy efficiency. The low-synchronization mode serves as an adaptive strategy that prioritizes functional demands at the expense of energy efficiency. Experimental data further confirm that during stress (e.g., mental arithmetic tasks), oxygen uptake increases while *HBC* decreases, accompanied by reduced energy efficiency. Moreover, dynamic changes in *HBC* strongly correlate with variations in cardiopulmonary energy efficiency, demonstrating a significant positive relationship. This further shows that dynamical changes synchronization level of HR and respiration can balance functional demands and energy efficiency.

This work advances our understanding of HR-respiration coupling by providing a rigorous theoretical proof for the physiological significance of in-phase synchronization. This contribution is distinct from previous work: First, it establishes a mechanistic link that complements the phenomenological findings of prior physiological experiments [67, 68]. Second, by employing analytical optimization conditions, our framework formally proves that a zero-phase difference is the optimal solution for energy efficiency, an insight not attainable through the numerical approaches used in previous theoretical models [50, 69]. It should be noted that the current theoretical framework is primarily predicated on the physiological parameters of healthy individuals in a non-exercise condition. Although the quantification of cardiopulmonary coupling has been extensively studied for applications in emotion classification, sleep monitoring, and disease surveillance, its specific quantification methods and physiological implications remain controversial. Our theoretical results demonstrate that HR-respiration synchronization maximizes cardiopulmonary energy efficiency, supporting the hypothesis that RespHRV serves as an intrinsic resting function of the cardiopulmonary system [67]. This also highlights the physiological significance of zero phase difference between HR and respiration, which was previously only demonstrated in experimental studies on dogs [68]. These findings further suggest that phase difference must be considered in mathematical methods for quantifying cardiopulmonary coupling.

Our experimental results show that the HR-respiration synchronization level operates along a dynamic continuum rather than exhibiting binary RespHRV/LF state transitions. The proposed *HBC* metric quantitatively captures this continuum, which show significant correlation with cardiopulmonary energy efficiency. This reveals an optimal regulatory strategy where the human body fine-tunes the RespHRV/LF ratio to balance energy conservation and functional demands. Notably, *HBC* provides a viable solution for noninvasive, continuous monitoring of cardiopulmonary efficiency, as its computation requires only HR and respiratory data.

The experimental results show that the energy saving caused by changes in the HR-respiration synchronization levles are of physiological significance. Classic literature [70] indicates that a typical healthy adult expends approximately 81.4 J/min of cardiac work under resting conditions [70]. In our experiments, the cardiac work during the stressful mental arithmetic task exceeds this value, whereas the four relaxation scenarios demonstrates reduced cardiac energy expenditure. This indicates that our experimental scenarios effectively induce the stress or more relaxed mood. On the other hand, it also shows that compared with the resting state, the cardiac energy consumption of the mental arithmetic task is approximately 7.47 J/min higher. This implies that if a person remains in a state of stress or working for 8 hours a day, the heart will consume approximately 3585.60 J more energy. On the other hand, relaxation induced through smoothing music and pleasant memories demonstrates greater cardiac work efficiency compared to ordinary rest, achieving more significant energy conservation. This indicates the importance of deep relaxation —such as that attained through meditation and mindfulness practices — for health maintenance and well-being.

It should be noted that since some physiological parameters cannot be directly measured noninvasively, our experimental results are obtained by combining measurable physiological quantities with a gas exchange dynamics model [49]. Although this introduces a potential source of discrepancy, we have undertaken multiple steps to ensure and demonstrate the model’s reliability. Firstly, the physiological quantities obtained by the model [49] are all within the normal physiological range for the healthy human, which include that the oxygen exchange efficiency in the lung, the volume of oxygen uptake and the lung capillary saturation of oxygen (*S*_*c*_*O*_2_). Mean values of these quantities for all volunteers calculated by the model are 4.2%, 233.1 ml and 97.4% respectively, which correspond to normal physiological values [71, 72]. Furthermore, we also utilize the experimental data of the finger blood oxygen saturation (*S*_*p*_*O*_2_) to estimate the blood oxygen partial pressure and derive the volume of oxygen uptake 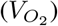 by *S*_*p*_*O*_2_, which is consistent with the results calculated by the model (shown in the Appendix F).

Consequently, while minor discrepancies with real data persist, as illustrated in Fig. 6 (which may stem from measurement noise or participant adherence), they do not undermine the model’s core utility or our conclusions. Future research incorporating advanced measurement technologies could further enhance the precision and comprehensiveness of physiological parameter quantification, thereby enabling more nuanced mechanistic interpretations. Futhermore, more sophisticated experimental protocols would help to investigate its broader implications: for example, adopting fully randomized scenario and item sequences to better control potential order effects, expanding the range of emotional or physiological scenarios tested, and including more diverse participant cohorts, etc.

Intriguingly, we identify analogous **bimodal coupling regulation in the pancreatic islet system**. In fact, our theoretical prove depends minimally on the details of cardiopulmonary coupling system, suggesting that the proposed bimodal optimization strategy may generalize to other oscillatory subsystems in living organisms. Experimental studies [73, 74] demonstrate that insulin (secreted by *β*-cells) and glucagon (secreted by *α*-cells) exhibit *in-phase* oscillations under normoglycemic conditions but switch to *anti-phase* oscillations during hyperglycemia (Fig. 8a-b). This can be formalized as an optimization problem based on *α*–*β* cell coupling:

**Fig. 8:**
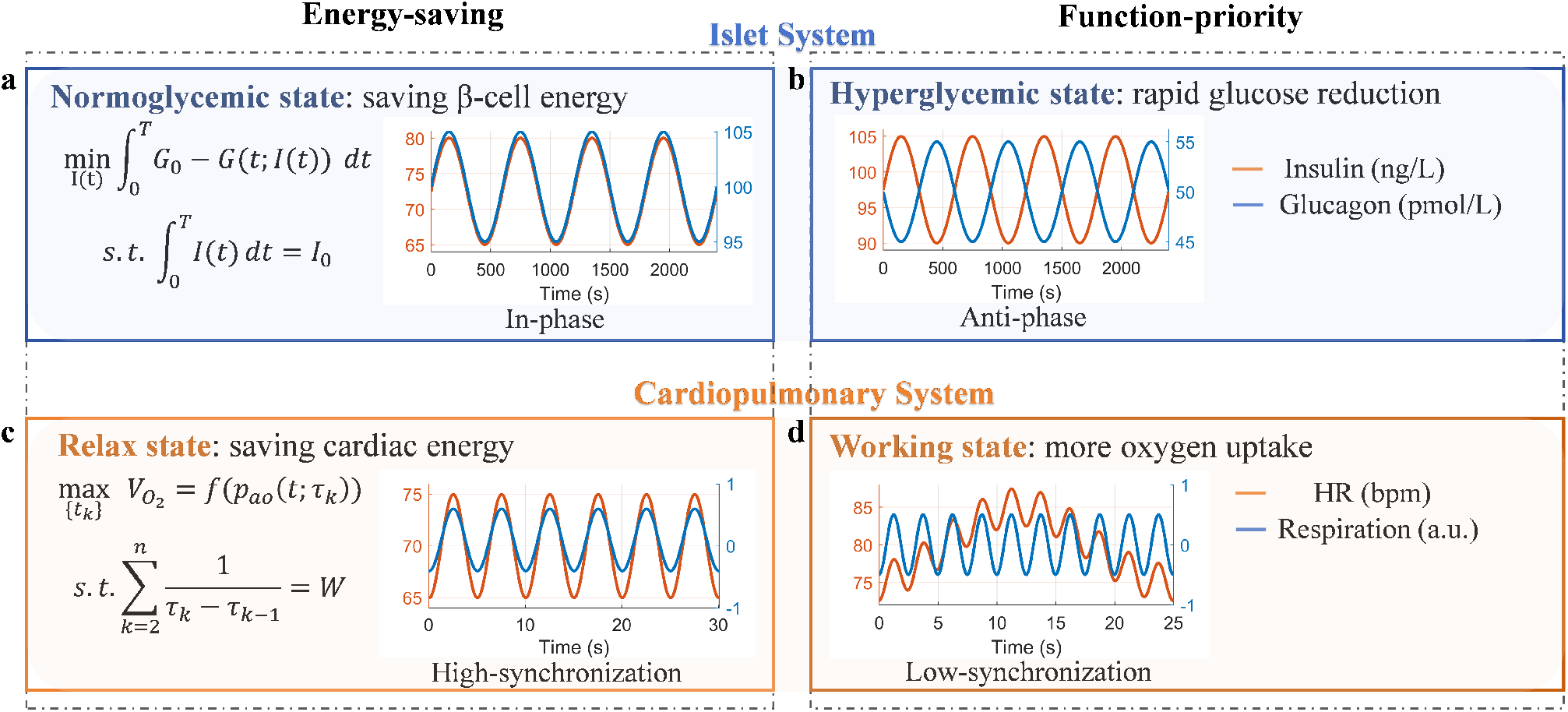
Bimodal coupling strategies in the Cardiopulmonary System and Islet System. **(a)** and **(b)** respectively show the oscillatory patterns of insulin (red) and glucagon (blue) in the pancreatic islet system under normoglycemic and hyperglycemic conditions. During normglycemia, their in-phase oscillations may minimize blood glucose fluctuations (maintaining glucose homeostasis) without increasing energy expenditure for *β*-cell insulin secretion. Under hyperglycemic conditions, insulin secretion increases and the two hormones transition to anti-phase oscillatory patterns to meet the functional requirement for rapid blood glucose reduction. **(c)** and **(d)** respectively show HR and respiration in the cardiopulmonary system during relaxed and stressed states. In the relaxed state, their in-phase synchronized oscillations maximize oxygen uptake with a restricted cardiac work. Under stress, this synchronization is disrupted, with the emergence of LF components in HRV. (a) and (c) correspond to the synchronized energy-saving mode within our proposed bimodal optimization framework, while (b) and (d) represent the desynchronized functional-priority mode.

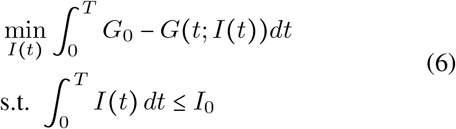

where *I* (*t*), *G* (*t*), *G*_0_, and *I*_0_ represent insulin concentration dynamics, blood glucose variation, baseline glucose level, and total insulin secretion from *β*-cells, respectively. The objective function quantifies the magnitude of blood glucose concentration fluctuations (with smaller values being preferable for maintaining glucose homeostasis). The constraint specifies the maximum total concentration of insulin secreted during time 0 − *T* by *β*-cells, representing the maximal energy expenditure during insulin secretion. This optimization framework formalizes how to allocate oscillatory insulin concentrations to minimize blood glucose variability while preserving *β*-cell energy expenditure. Under hypoglycemic conditions, the glucose concentration *G* (*t*) approximates the basal level *G*_0_. This formulation appears to align with the HR-respiratory coupling optimal problem (as shown in Fig. 8a) and shares conceptual similarities with the *water-filling principle* from communication theory: given a maximal total insulin output, temporal allocation of secretion may optimize glycemic stability. The optimal solution under normoglycemia could involve *in-phase α*–*β* oscillations, potentially maximizing energy efficiency. During hyperglycemia, preliminary observations [75] indicate that functional demand for rapid glucose reduction may require increased insulin secretion, which appears to disrupt synchronized *α*–*β* oscillations (as shown in Fig. 8b) - a pattern resembling our cardiopulmonary observations. However, more rigorous theoretical exploration of this optimization problem requires an established and physiologically plausible model of *α*–*β* cell interactions. Therefore, we cannot provide the stringent theoretical proofs.

The bimodal optimization paradigm opens a new perspective for understanding intelligent regulation in biological systems. In systems biology, this paradigm provides a novel approach to investigate oscillatory coupling, as demonstrated by the HR-respiration coupling in the cardiopulmonary system, with preliminary evidence also suggesting its applicability to the pancreatic islet system. For health preservation, our findings emphasize the importance of work-rest balance, as continuous high intensity work, chronic stress and excessive anxiety will deplete cardiac energy reserves. From the interdisciplinary perspective, the integration of eastern “Yin-Yang” philosophy and “Heart-Breath Interdependence” Daoist cultivation principle with western science (specifically, systems physiology, dynamical systems modeling, and optimization theories from engineering), offers unique insights into optimization strategies for complex oscillatory networks.

## Methods

### Proof of Theorem 1

Theorem 1 can be proved in three steps:

Step1 - Derive an explicit expression of the objective function.

First, consider the oxygen uptake on each R-R interval:

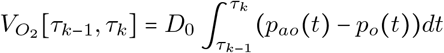

for *k =* 1, 2, …, *n*. Direct measurement of *p*_*ao*_ (*t*) and *p*_*o*_ (*t*) is generally difficult and invasive, but they can be obtained via a gas exchange dynamic model [49] as the following:

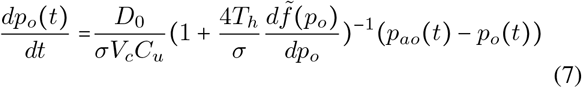

where the function 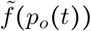 represents the oxygen-hemoglobin dissociation curve:

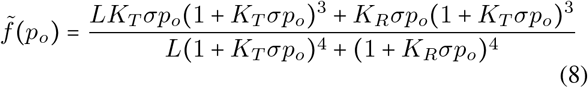

which is a monotonously increasing function, i.e., 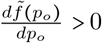, implying a strict rise in the percentage of hemoglobin that becomes oxygen-bound as the partial pressure of oxygen in the blood increases [51]. During each RR interval [*τ*_*k*−1_, *τ*_*k*_], the partial pressure of oxygen *<u>p</u>*_*o*_ *<u>t</u>* demonstrates a consistent increasing trend, implying 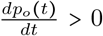 throughout this period. Consequently, 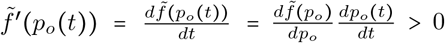, for *t* ∈ [*τ*_*k*−1_, *τ*_*k*_]. The constants *σ* =1.4 × 10^−6^*L*^−1^*mmHg*^−1^, *V*_*c*_ = 0.07*L, C*_*u*_ = 25.426*Lmol*^−1^, and *T*_*h*_ = 2 × 10^−3^*molL*^−1^ are oxygen solubility in blood plasma, capillary volume, unit conversion factor, capillary haemoglobin concentration, respectively [49].

Then, the oxygen uptake on each R-R interval can be calculated as the following:

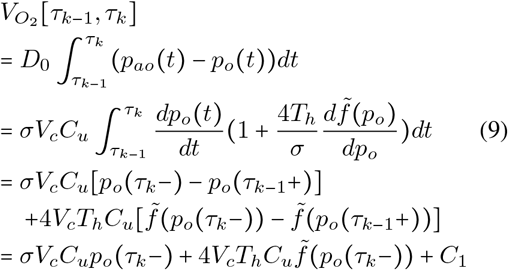

where (9) follows from (7), and *C*_1_ is some constant since immediately after each heartbeat *p*_*o*_(*τ*_*k*−1_+) can be assumed to be the same. According to the basic physiology [51], when blood has moved a third of the distance through the capillary, its oxygen partial pressure rises to a level that is approximately consistent with the oxygen partial pressure in the alveoli, which means that *p*_*o*_ (*τ*_*k*_^−^) ≈ *p*_*oa*_ (*τ*_*k*_). Therefore, noting that 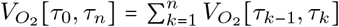, the objective function (1) can be replaced by:

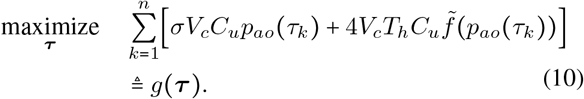

Step2 - Apply the KKT conditions

The optimization problem (10) subject to (2) can be solved using a method analogous to the water-filling principle [56, 57]. First, construct the Lagrange function as follows:

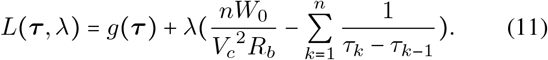

Based on the KKT conditions, the optimal heartbeat sequence 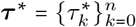 should meet the following criteria:

1. Primal Feasibility:

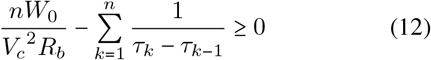
2. Dual Feasibility:

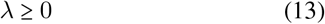
3. Complementary Slackness:

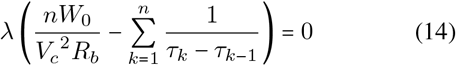
4. Stationarity: For *k* = 1, 2, …, *n* − 1, where

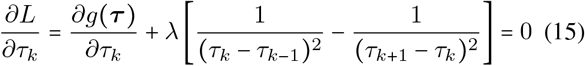

where

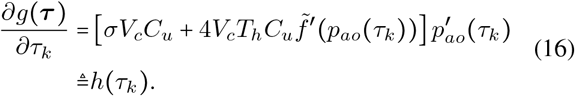

Because 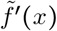 is always positive as noted earlier, we have 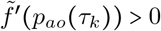. Therefore, the sign of *h*(*τ*_*k*_) is determined by the sign of 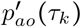.

Step3 - Solve for the optimal heartbeat timings

Since *p*_*ao*_(*t*) is not constant throughout the respiratory cycle, which guarantees the existence of at least one *i* ∈ {1, …, *n* − 1} such that 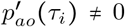 ≠ 0, it follows from (16) that *h*(*τ*_*i*_) ≠ 0 for such *i*. Therefore, we must have *λ* ≠ 0 by (15). Hence by (13), we have the solution:

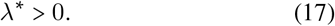

Hence by (14),

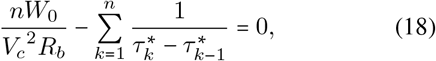

which means the optimum is achieved precisely when the total cardiac work equals *nW*_0_.

Combining (15) and (18), we obtain Theorem 1.

**Proof of Corollary 1.1**

It has been shown in [58–62] that when the amount of inhaled gas is sufficient (non-hypoxic), the temporal derivative of alveolar oxygen partial pressure exhibits the following behavior during the interval between the second and penultimate heartbeats within one respiratory cycle:

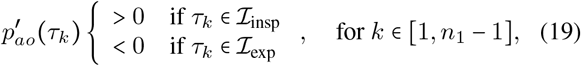

where *τ*_1_ and 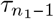 denote the timings of the second and penultimate heartbeats during one respiratory cycle, respectively. It should be noted that due to the presence of alveolar dead space (represented by the model parameter *V*_*D*_, more detalis in Appendix A), *p*_*ao*_ (*t*) exhibits a brief initial decrease (lasting approximately 0.5 seconds [58]) at the start of inspiration before establishing the characteristic pattern of increase during inspiration and decrease during expiration. Since our subsequent proofs only require the behavior of *p*_*ao*_(*t*) after the second heartbeat, we restrict the index *k* in *τ*_*k*_ accordingly to exclude this initial transient phase. Therefore, by (16),

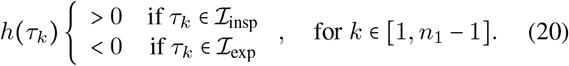

Next, Corollary 1.1 easily follows from (3) and (20)-(17).

### Numerical solution of the HR-respiratory coupling mode with optimal energy efficiency

The numerical solution to the optimization problem is obtained using the simulated annealing algorithm. The initial temperature is set to 100, and decreases continuously according to the formula *θ*_*k*+1_ = *θ*_*k*_ × 0.95. When the temperature drops to 10^−4^, the algorithm switches to the gradient descent method with learning rate as 0.1. The convergence condition is set as the gradient magnitude being less than 10^−5^ or reaching the maximum number of iterations of 5000. To optimize the observation of the phase relationship between peak heart rate timing and respiration, the constrained cardiac work (*W*_0_) is determined for each specified respiratory cycle duration *T*. For each *T* value (3, 4, 5, 6, 7, 8, 9, 10), *W*_0_ is set to its maximum permissible value, subject to two critical constraints: (1) the mean heart rate remains below 90 bpm, a physiologically appropriate threshold aligned with empirical observations from experimental data; (2) the total heartbeat count is an even number (thereby ensuring an odd RR interval count). The resulting *W*_0_ values are 1.00, 1.25, 1.41, 1.16, 1.28, 1.36, 1.22, 1.49, respectively. The time duration [*τ*_0_, *τ*_*n*_] are set as 10 complete respiratory cycles. After obtaining the optimal solution, the average of the optimal heart rates over respiratory cycles is calculated and plotted in Fig. 2b.

### Calculation of cardiac work

The heart pumps out a volume *V*_*c*_ of blood with pressure *P* during each R-R interval (*RR*_*i*_) as proposed in [50], denoting the blood flow resistance as *R*_*b*_:

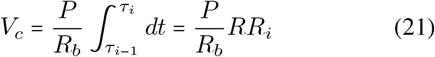

Therefore, the cardiac work *W* (*J*) per heart beat can be computed by:

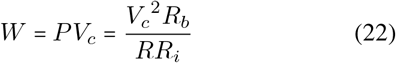

The heart stroke volume *V*_*c*_ and *R*_*b*_ are set as 70 *ml* and 1119 *mmHg s L*^−1^ (mean in the physiological range [76– 78]) respectively.

### Calculation of oxygen uptake and carbon dioxide output

The computation of pulmonary oxygen uptake 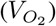 and carbon dioxide output 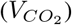 involve alveolar oxygen partial pressure (*p*_*ao*_), alveolar carbon dioxide partial pressure (*p*_*ac*_), blood oxygen partial pressure (*p*_*o*_), and blood carbon dioxide partial pressure (*p*_*c*_). These parameters cannot be directly measured non-invasively and continuously. Therefore, we employ continuously measurable physiological signals (HR and respiration) combined with a gas exchange dynamic model [49] to calculate 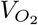 and 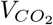 . Appendix A details the gas exchange dynamic model used and the methodology for calculating oxygen uptake by incorporating actual heart rate and respiratory data into the model.

### Experimental data

The database of 65 healthy volunteers aged 27.4 ± 8.9 years (31 males and 34 females) are recorded. The HR, respiration and fingertip oxygen saturation are measured synchronously by portable Embletta MPR PG device (Natus Medical Inc., San Carlos, CA, USA), and the frequency of output data are 5hz, 5hz and 3hz respectively. Continuous eletrocardiography (ECG) data is acquired using 3-channel recordings with the sampling rate of 250 Hz, and the respiratory data is acquired by the belt-type sensor placed on the chest and abdomen with the sampling rate of 100 Hz.

Ethics Committee at the Department of Psychology, Tsinghua University approved the whole experimental setup, and all volunteers gave their written consents before the experiment. Alcohol and vigorous exercise are forbidden for 24 hours and an hour before the experiment. The experiment was conducted between 7 and 9 p.m. every night and lasted for about an hour. At the beginning of the experiment, volunteers were instructed to breathe deeply and slowly to the rhythm of the metronome, which allowed volunteers to get used to the site and equipment and to eliminate effects of potential stress.

The experiment includes five scenarios to induce relax or stress state of the voluteers. and the detailed experimental process is as follows:

1. Scenario 1 - arithmetic task: volunteers have to count down from 1000 in steps of 17 for one minute, and are told to present their calculation results at the end of this item. They are considered to be under stress in this stage.
2. Scenario 2 - listening to sounds: volunteers listen to (i) the blue noise, (ii) the smooth piano music, (iii) quarrel audio with their eyes closed, and each lasts for 90 s.
3. Scenario 3 - emotional evocations: volunteers are instructed to recall (i) positive and relaxing moments, (ii) negative and annoying moments by picture and voice guidance. Each item lasts for 150 s and the data is used for analysis in the last 60 s, which is the most intense stage of volunteers’ emotions.
4. Scenario 4 - smelling odors: volunteers are requested to smell (i) essential oils, (ii) toilet clean spirit, (iii) Chinese herb and they do not know what they will smell before each item. Each odor lasts for 90 s.
5. Scenario 5 - watching videos: volunteers watch videos without sounds about (i) lovely flower or pet, (ii) scary spiders. Each video lasts for 60 s.

In experimental scenarios 2-5, listening to piano music, recalling positive moment, smelling essential oil, and watching lovely flower or pet video are implemented to induce a more relaxed emotional state. A resting stage of 45 s is set between each item, during which volunteers were instructed to take deep breaths. In the actual implementation of the experimental protocol, the sequence of scenarios and items was not identical for all participants. Specifically, for scenario order: 15 volunteers followed the 1-2-3-4-5 order, 14 volunteers completed scenarios in the 1-2-5-3-4 order, and 10 volunteers followed the 1-2-3-5-4 order, with other sequences assigned to a relatively smaller number of volunteers. Additionally, the order of experimental items within each scenario was adjusted; for instance, in scenario 3, both positive-negative and negative-positive item presentation orders were implemented.

After the experiment, visual analogue scale (VAS) is used to get feedback on how relaxed and stressed volunteers feel during each item. In addition, for the emotional evocation scenario, volunteers are asked to report whether they really introduced the corresponding emotion, so as to delete some unqualified data.

### Validation of the gas exchange dynamic model by experimentally measured fingertip blood oxygen

Since our conclusions depend on the gas exchange dynamic model [49], validating its effectiveness and reliability is essential. Appendix F presents the detailed validation strategies and their corresponding results. Specifically, we validate the model outputs using measured fingertip blood oxygen levels. The validation process involves two key analyses: (1) comparative assessment between the real finger-tip blood oxygen (*S*_*p*_*O*_2_) variations and the model-derived arterial blood oxygen (*S*_*a*_*O*_2_) fluctuations, and (2) investigation of whether oxygen uptake estimates based on fingertip measurements exhibits consistent conclusion with model-calculated oxygen uptake - regarding whether the oxygen uptake per unit cardiac work (energy efficiency) still maintains a positive correlation with HR-respiration synchronization level (*HBC*). Results show that: (1) the mean value of the difference bewteen *S*_*a*_*O*_2_ and *S*_*p*_*O*_2_ is only 2.12% (Appendix Fig. S 3a). *S*_*a*_*O*_2_ and *S*_*p*_*O*_2_ are positive correlated with *r* = 0.69 and *p* < 0.001 (Appendix Fig. S 3b); (2) the *HBC* also demonstrates significant positive correlation with the *S*_*p*_*O*_2_-estimated oxygen uptake with *r* = 0.69 and *p* <0.01. (Appendix Fig. S 4b). These results affirmatively confirm both propositions, thereby validating the reliability of the model.

### Analysis and statistics

The normality of the paired differences is assessed using the Shapiro-Wilk test. For comparisons where the paired differences do not deviate significantly from normality, two-sided paired t-tests are employed. These results are reported with the mean difference, its 95% confidence interval (CI), the t-statistic, and the associated p-value. In cases where the normality assumption is violated, two-sided Wilcoxon signed-rank tests are performed. For these tests, we report the V statistic, p-value, and the Hodges-Lehmann estimate of the location shift with its 95% CI. Effect sizes for paired comparisons are calculated as Cohen’s d, defined as the mean of the paired differences divided by their standard deviation. This is computed as:

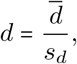

where for each pair i,

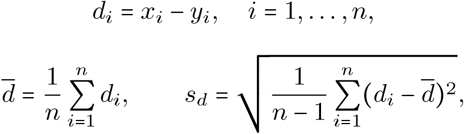

Here *d* is the sample mean of the differences, *s*_*d*_ is their sample standard deviation, and *n* is the number of pairs. For correlation analyses, the normality of each variable is examined separately using the Shapiro-Wilk test. If both variables are normally distributed, Pearson’s correlation coefficient is computed along with its p-value. Otherwise, Spearman’s rank correlation and its p-value are reported.

## Author contributions

J.H.: Conceived the research framework, formulated core hypotheses and conclusions. J.Z.: Executed the experiments and analyzed the experimental data, conducted simulations and mathematical proofs. L-L.X.: Supervised the mathematical proofs, provided the key proof idea and method. J.Z. and J.H.: Writed the original draft. All authors contributed to critical review, editing, and final approval of the manuscript.

